## Supplementary Figures and Tables for "FTO intronic SNP strongly influences human neck adipocyte browning determined by tissue and PPARγ specific regulation: a transcriptome analysis"

**Supplementary Figure 1.**

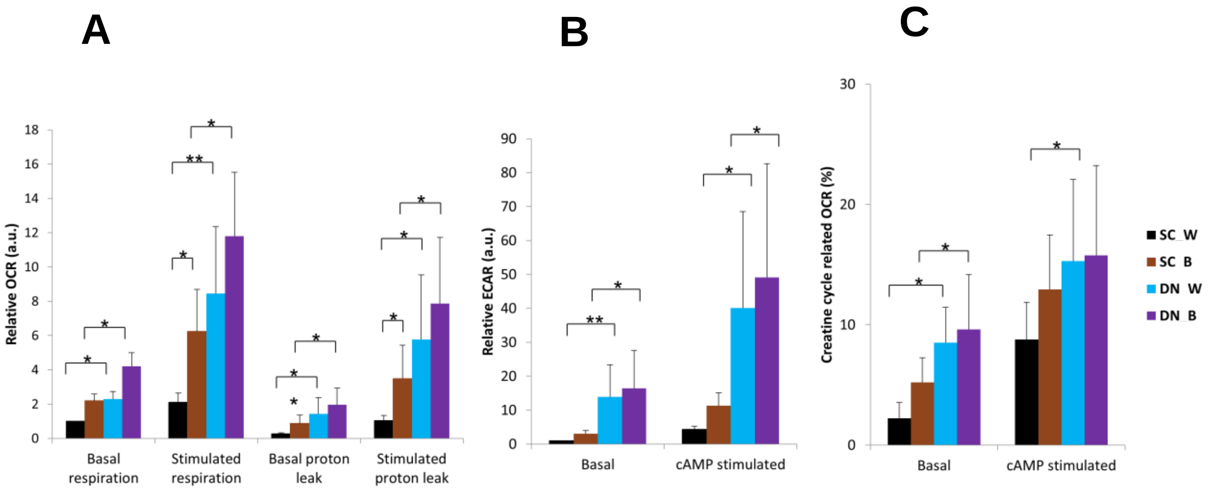


**Supplementary Figure 1. Functional features of the primary SC and DN adipocytes differentiated for two weeks to white or brown. (A) Basal, cAMP stimulated and oligomycin inhibited oxygen consumption (OC) levels** (as compared to basal OCR of SC white adipocytes) n=4. **(B) Basal and cAMP-stimulated extracellular acidification levels** (as compared to basal ECAR of SC white adipocytes) n=4. **(C) The proportion of creatine kinase futile cycle related OC at basal and cAMP-stimulated** n=4. SC: Subcutaneous, DN: Deep-neck, W: white differentiation protocol, B: brown differentiation protocol. Statistics: In comparison of two groups two-tailed paired Student’s t-test was used. *$p<0.05$. In multi-factor comparision we used two-way ANOVA and post hoc Tukey’s test. *$p<0.05$. **$p<0.01$. n=4

### Supplementary Figure 2.

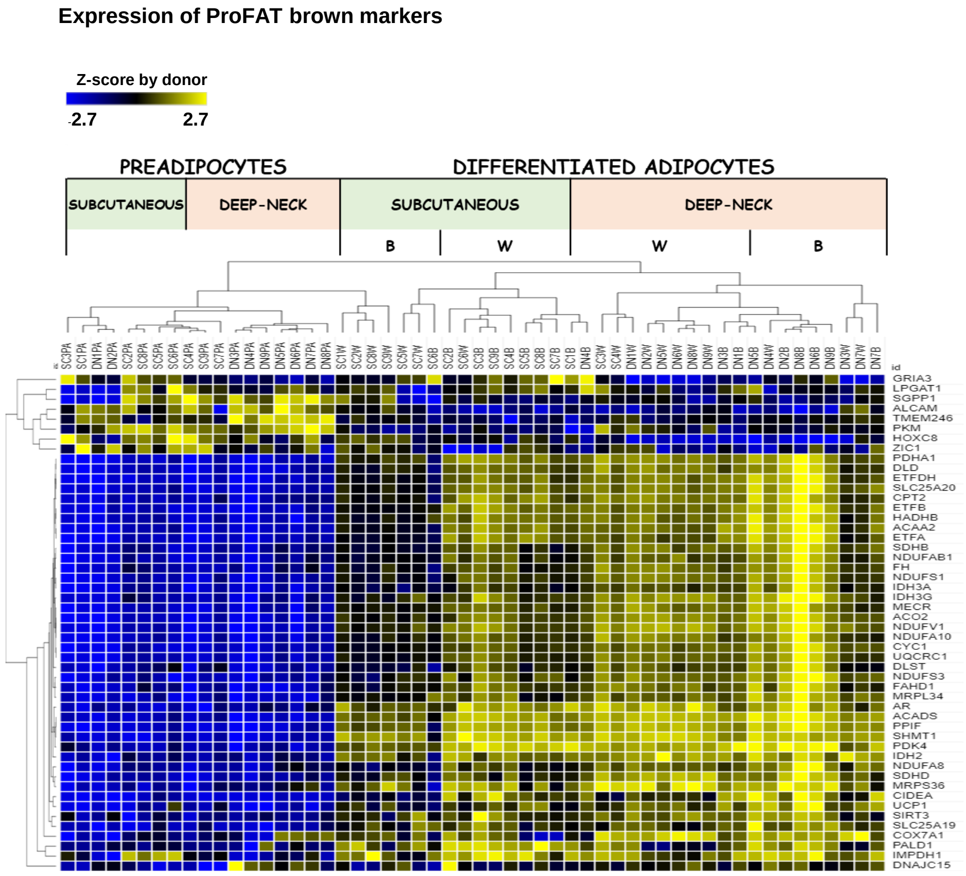


**Supplementary Figure 2. Expression profile of ProFAT marker genes.** Heat map shows the expression profile of browning (44) and white (6) characteristic marker genes from ProFAT database in SC and DN adipose progenitors and differentiated samples.

**Supplementary Figure 3.**

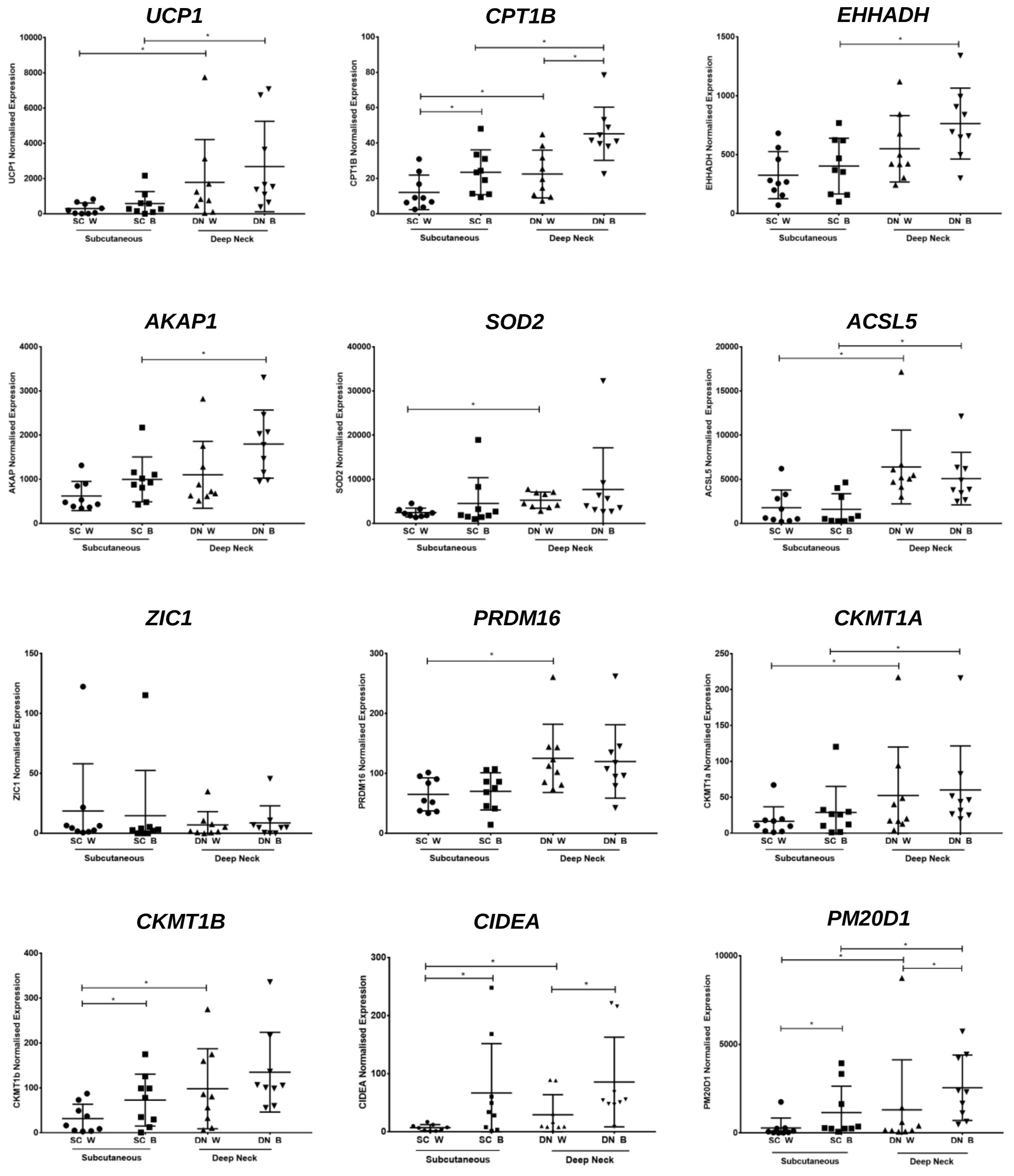


**Supplementary Figure 3. Gene-expression of key browning marker genes based on RNAseq data**. SC: Subcutaneous DN: Deep-neck. W: white differentiation protocol; B: brown differentiation protocol; Statistics: Deseq R data. *p<0.05. n=9 donors. 4 samples/donor.

**Supplementary Figure 4.**

**A**

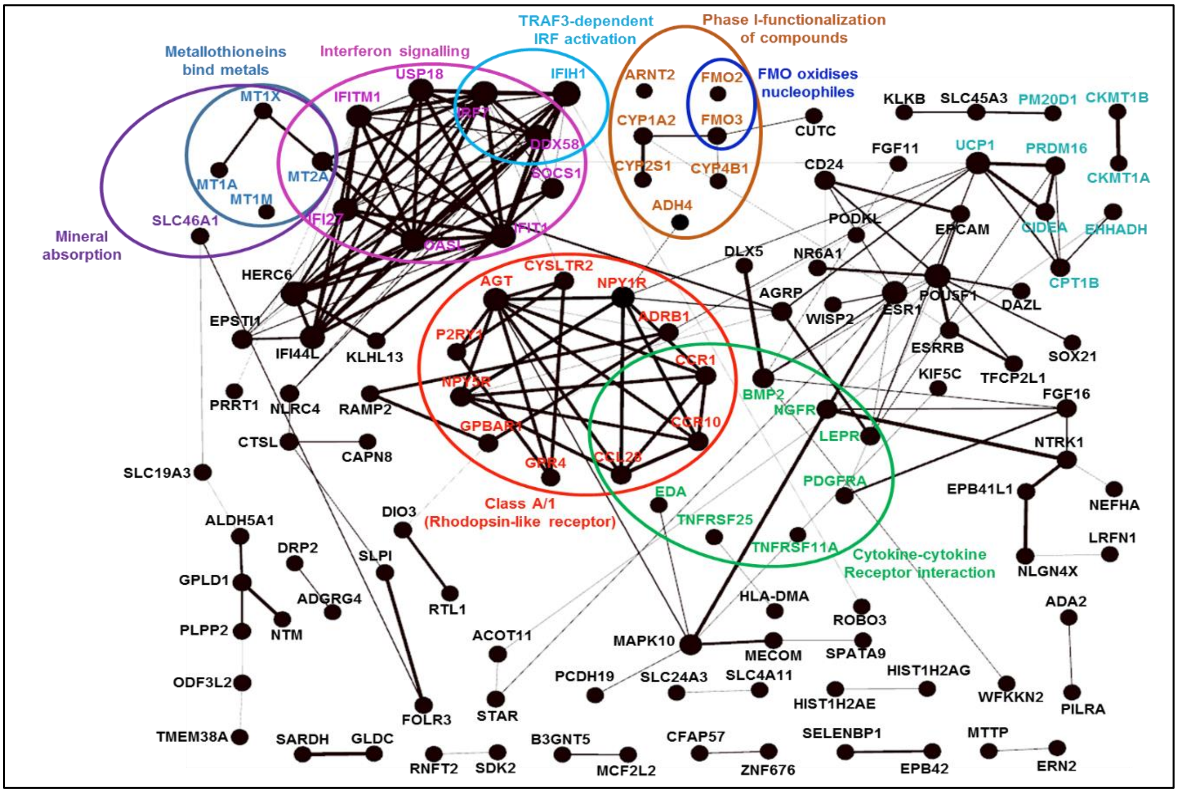


**B**

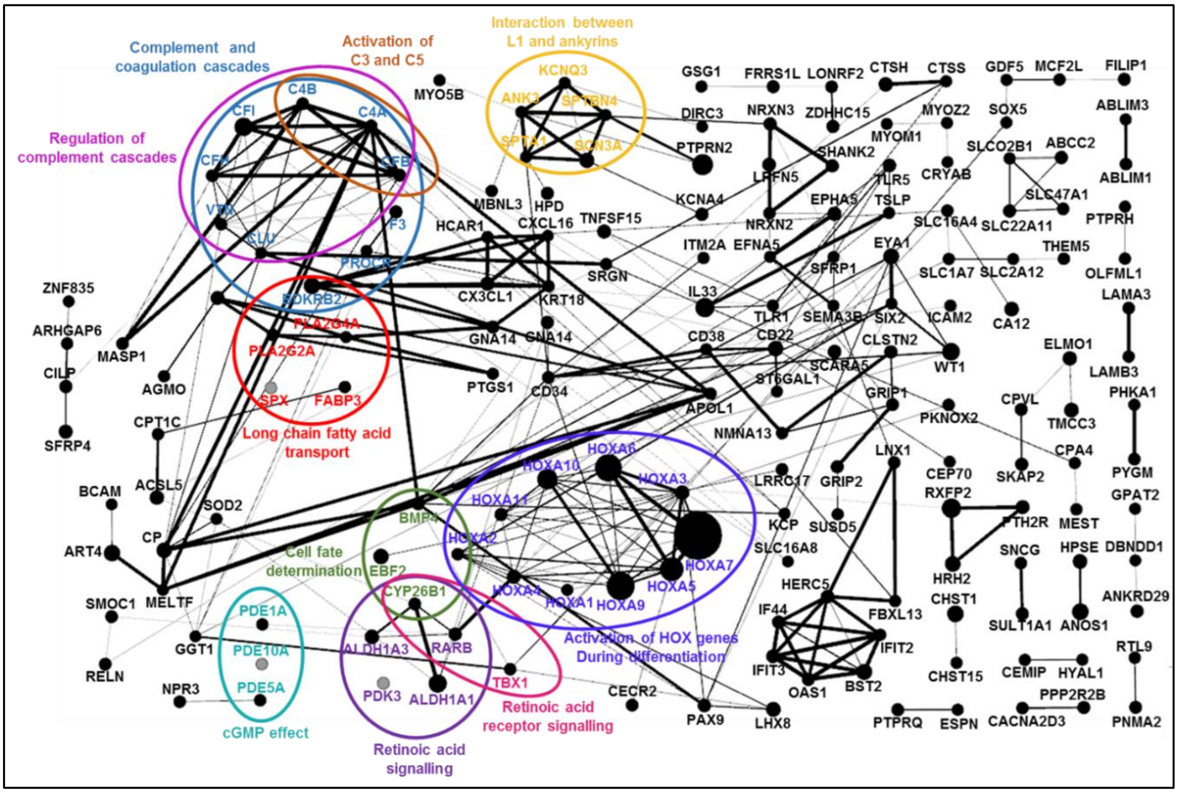

C

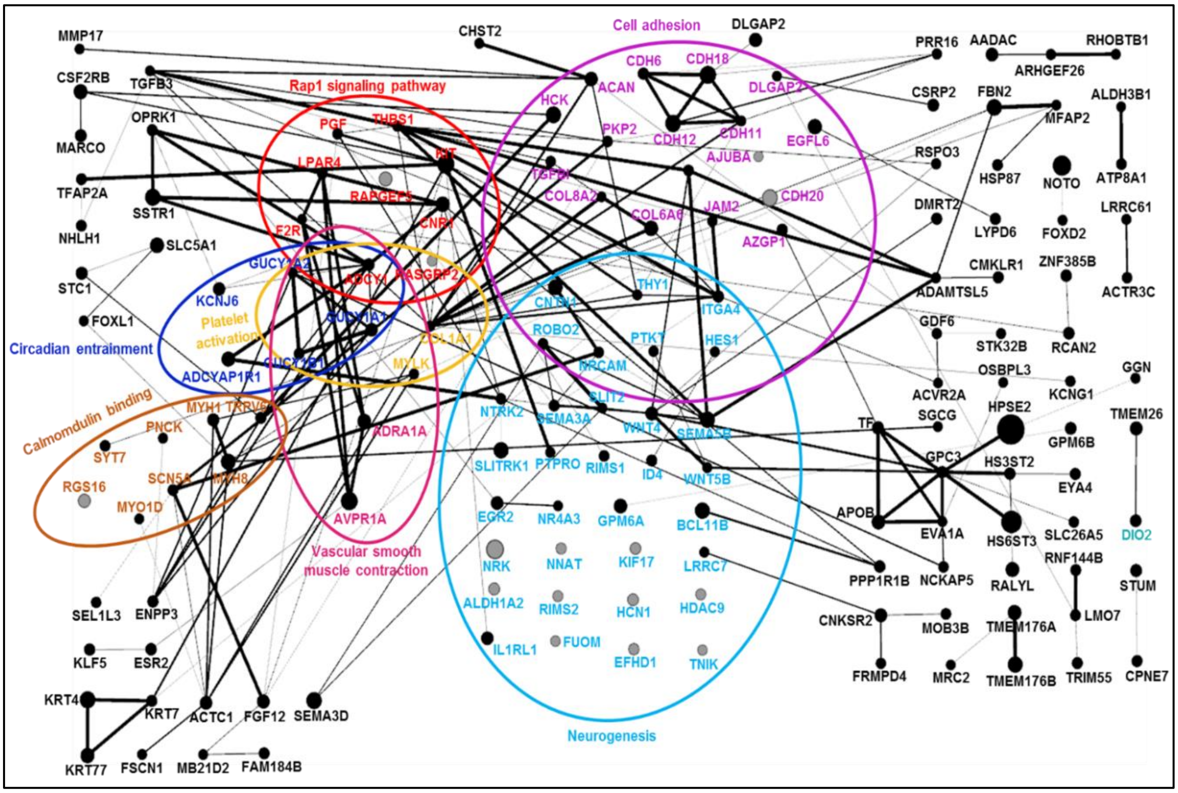

Rap1 signaling pathway
Cell adhesion
Circadian entrainment
Platelet activation
Calmodulin binding
Vascular smooth muscle contraction
Neurogenesis
MMP17
CHST2
DLGAP2
PRR16
AADAC
RHOBTB1
ARHGEF26
CSF2RB
TGFB3
CDH6
CDH18
DLGAP2
ACAN
HCK
CSRP2
FBN2
MFAP2
ALDH3B1
OPRK1
PGF
THBS1
PKP2
CDH12
CDH11
EGFL6
MARCO
LPAR4
KIT
AJUBA
RSPO3
HSP87
NOTO
ATP8A1
TFAP2A
SSTR1
RAPGEF3
CNR1
COL8A2
COL6A6
JAM2
CDH20
DMRT2
LRRC61
NHLH1
F2R
AZGP1
LYPD6
FOXD2
SLC5A1
GUCY1A2
ADCY1
RASGRP2
CMKLR1
ZNF385B
STC1
KCNJ6
CNTN1
THY1
ITGA4
ADAMTSL5
ACTR3C
FOXL1
GUCY1A1
COL1A1
ROBO2
PTK7
HES1
GDF6
STK32B
RCAN2
ADCYAP1R1
GUCY1B1
MYLK
NRCAM
OSBPL3
GGN
SLIT2
ACVR2A
KCNG1
MYH1
TRPV6
NTRK2
SEMA3A
WNT4
SEMA3B
SGCG
HPSE2
TMEM26
PNCK
TF
SYT7
ADRA1A
SLITRK1
PTPRO
RIMS1
ID4
WNT5B
GPC3
HS3ST2
GPM6B
RGS16
SCN5A
MYH8
EYA4
MYO1D
AVPR1A
EGR2
NR4A3
GPM6A
BCL11B
APOB
EVA1A
HS6ST3
SLC26A5
DIO2
NRK
NNAT
KIF17
LRRC7
RNF144B
PPP1R1B
NCKAP5
RALYL
STUM
ALDH1A2
RIMS2
HCN1
HDAC9
SEL1L3
ENPP3
CNKSR2
MOB3B
TMEM176A
LMO7
IL1RL1
FUOM
EFHD1
TNIK
KLF5
ESR2
FRMPD4
MRC2
TMEM176B
TRIM55
CPNE7
KRT4
KRT7
ACTC1
FGF12
SEMA3D
KRT77
FSCN1
MB21D2
FAM184B


D

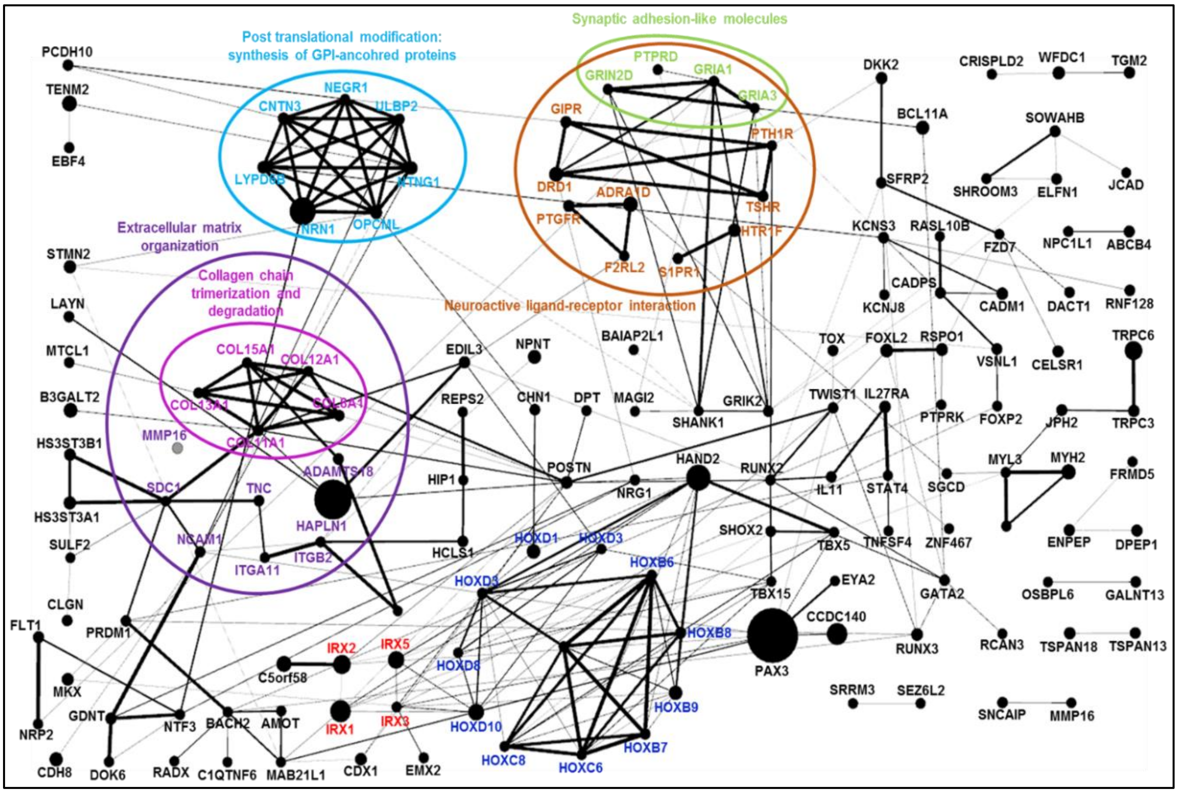

Post translational modification: synthesis of GPI-ancohred proteins
Synaptic adhesion-like molecules
Extracellular matrix organization
Collagen chain trimerization and degradation
Neuroactive ligand-receptor interaction
PCDH10
TENM2
EBF4
NEGR1
CNTN3
ULBP2
LYPD6B
NTNG1
NRN1
OPCML
PTPRD
GRIN2D
GRIA1
GRIA3
GIPR
PTH1R
DRD1
ADRA1D
TSHR
PTGFR
HTR1F
F2RL2
S1PR1
DKK2
BCL11A
SFRP2
CRISPLD2
WFDC1
TGM2
SOWAHB
SHROOM3
ELFN1
JCAD
KCNS3
RASL10B
FZD7
NPC1L1
ABCB4
STMN2
CADPS
KCNJ8
CADM1
DACT1
RNF128
LAYN
TOX
FOXL2
RSPO1
VSNL1
CELSR1
TRPC6
MTCL1
COL15A1
COL12A1
COL13A1
COL8A1
COL11A1
EDIL3
NPNT
BAIAP2L1
B3GALT2
REPS2
CHN1
DPT
MAGI2
SHANK1
GRIK2
TWIST1
IL27RA
PTPRK
FOXP2
JPH2
TRPC3
MMP16
ADAMTS18
HS3ST3B1
HS3ST3A1
SDC1
TNC
HAPLN1
HIP1
POSTN
NRG1
HAND2
RUNX2
IL11
STAT4
SGCD
MYL3
MYH2
FRMD5
SULF2
NCAM1
ITGA11
ITGB2
HCLS1
HOXD1
HOXD3
SHOX2
TBX5
TNFSF4
ZNF467
ENPEP
DPEP1
HOXB6
HOXD4
TBX15
EYA2
GATA2
OSBPL6
GALNT13
CLGN
CCDC140
FLT1
PRDM1
HOXB8
PAX3
RUNX3
RCAN3
TSPAN18
TSPAN13
IRX2
IRX5
HOXD8
MKX
C5orf58
SRRM3
SEZ6L2
HOXB9
SNCAIP
MMP16
GDNT
NTF3
BACH2
AMOT
IRX1
IRX3
HOXD10
NRP2
HOXB7
HOXC8
HOXC6
CDH8
DOK6
RADX
C1QTNF6
MAB21L1
CDX1
EMX2

**Supplementary Figure 4. Interactome maps of group 1-4 genes (A) Interactome map of group 1 genes. (B) Interactome map of group 2 genes (C) Interactome map of group 3 genes (D) Interactome map of group 4 genes.** The interaction network was determined by STRING (https://string-db.org) and constructed by using Gephi 0.9.2 (https://gephi.org). Size of the nodes reflect the fold change of DN/SC (**A and B**) and SC/DN (**C and D**). Edges represent protein-protein interactions and the size of the edges indicate the confidence of interaction or the strenght of data support. The clustering was performed based on reactome (reactome.org) and KEGG Mapper (www.kegg.jp)

#### Supplementary Figure 5.

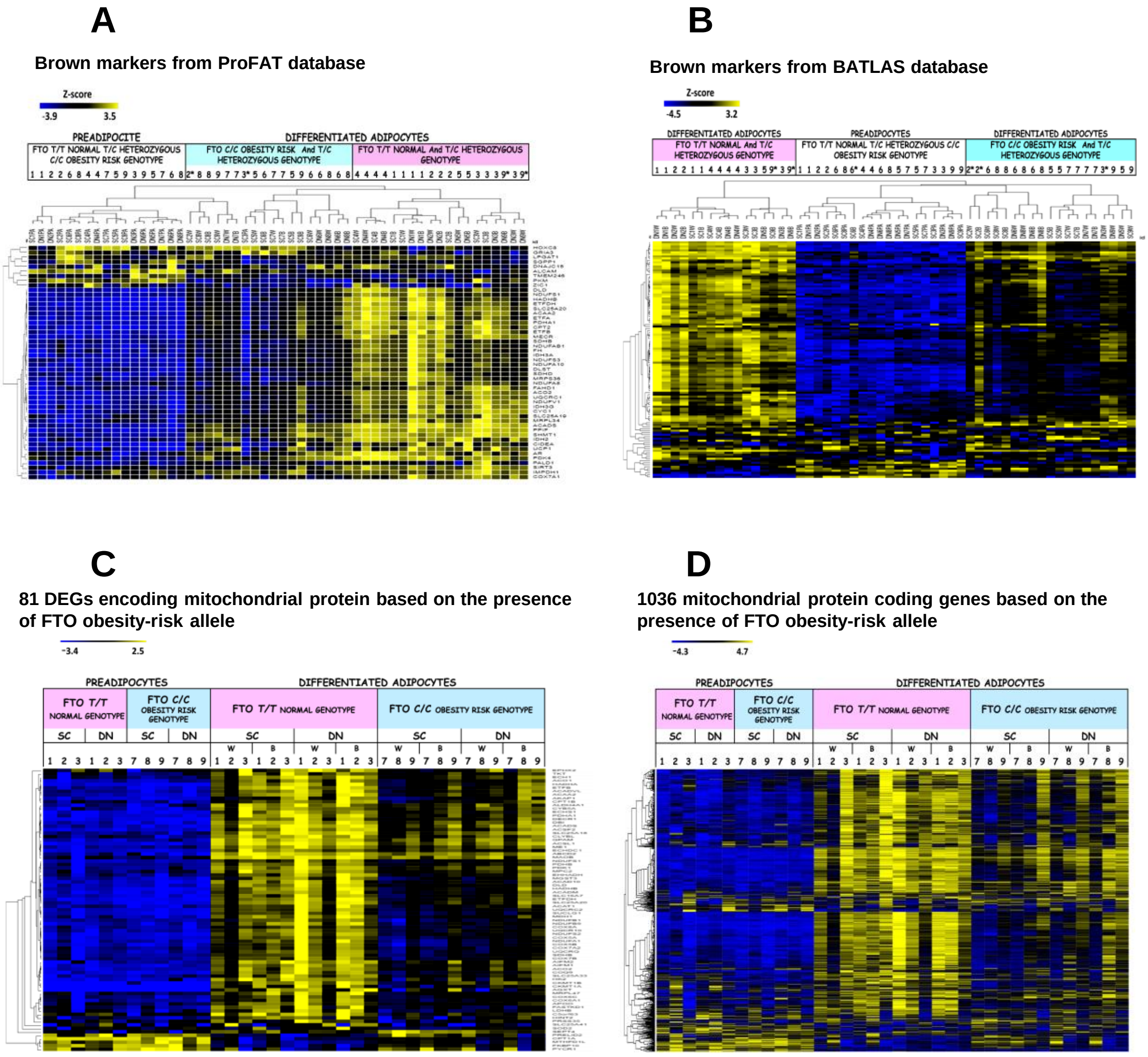


**Supplementary Figure 5. FTO obesity-risk allele presence based analyses of gene-expression profiles. (A) Expression profile of ProFAT marker genes**. Heatmap shows the expression profile of browning (44) and white (6) characteristic marker genes from ProFAT database in SC and DN adipose progenitors and differentiated samples; samples was hierarchically clustered based on pearson correlation. n=9 **(B) Expression profile of adipocyte specific BATLAS marker genes.** Heatmap shows the expression profile of browning (98) and white (21) characteristic marker genes from BATLAS database in SC and DN adipose progenitors and differentiated samples, samples was hierarchically clustered based on pearson correlation. To indentify donor differences z-score was calculated by considering all donors. n=9. **(C) Expression profile of the 81 genes encoding mitochondrial proteins**, which expressed differently according FTO obesity-risk allele presence. **(D) Expression profiles of mitochondrial proteins encoding genes based on Human MitoCarta 2.0 database.** Heat map shows the expression profiles of 1038 genes encoding mitochondrial proteins based on the presence of the *FTO* obesity-risk allele C/C in SC and DN adipose progenitors and differentiated samples. z-score was calculated by all samples to identify donor differences. n = 6 donors. SC Subcutaneous; DN: Deep-neck; W white differentiation protocol; B brown differentiation protocol. Numbers 1-9 represents donors, * outlier from the cluster.

#### Supplementary Figure 6.

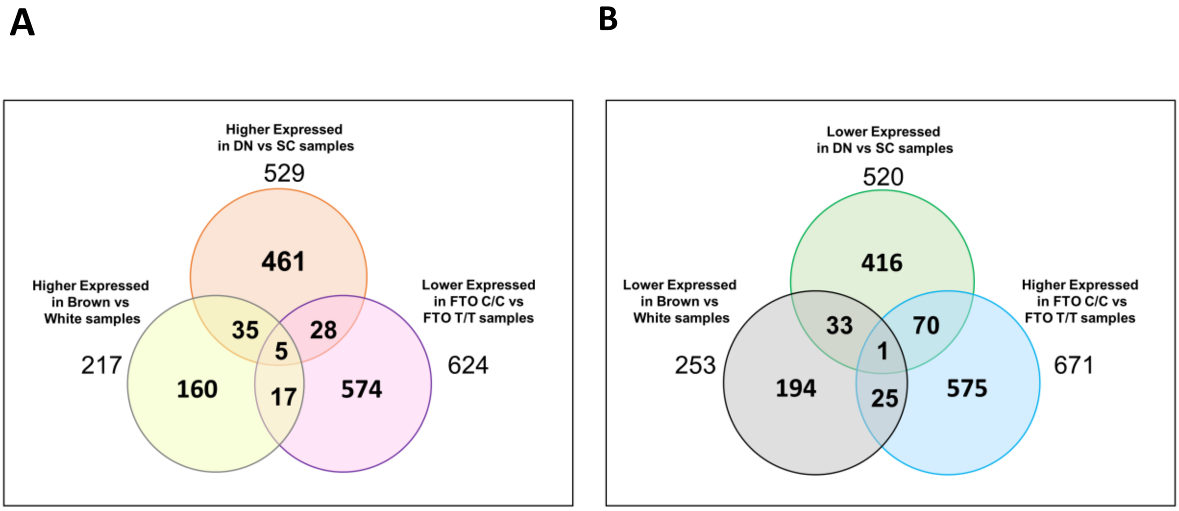


**Supplementary Figure 6. Number of differentially expressed genes in differentiated adipocytes based on tissue origin, differentiation protocol and the presence of the FTO obesity-risk allele. (A)** Venn diagram shows significantly higher expressed genes number in samples expected higher thermogenic activity, and among of these genes several were lower expressed in FTO obesity-risk genotype samples **(B)** shows significantly lower expressed genes number in samples expected higher thermogenic activity, and among of these genes many expressed higher in FTO obesity-risk genotype samples. SC: subcutaneous; DN: deep-neck.

**Supplementary Table 1.**

| Indicated cell type or component | CD markers | Percentage of positive cells (SC PA) | Percentage of positive cells (DN PA) |
|---|---|---|---|
| | *CD34* | 0.01±0.01 | 56.31±48.93 |
| | *CD45* | 0.00±0.00 | 0.00±0.00 |
| Hematopoietic/ Monocyte markers | *CD47* | 96.24±1.36 | 94.36±1.72 |
| | *CD338 (ABCG2)* | 27.32±1.36 | 35.61±32.46 |
| | *HLA-DR* | 30.22±52.34 | 28.46±49.29 |
| Endothelial markers | *CD31 (PECAM)* | 0.00±0.00 | 3.43±5.94 |
| | *CD54 (ICAM-1)* | 37.94±39.11 | 32.79±56.73 |
| | *CD73* | 96.04±1.84 | 94.49±5.59 |
| MSC/Fibroblast markers | *CD90 (Thy-1)* | 94.62±1.20 | 87.78±6.23 |
| | *CD105 (Endoglin)* | 56.81±49.23 | 20.53±35.56 |
| | *CD29 (Integrin β1)* | 87.93±15.62 | 97.27±3.23 |
| | *CD36* | 76.40±4.32 | 79.22±7.78 |
| | *CD44 (H-CAM.Hermes)* | 85.31±11.82 | 60.29±48.31 |
| Integrins and CAMs | *CD49a (Integrin α1)* | 50.50±45.76 | 68.26±4.89 |
| | *CD49d (Integrin α4)* | 27.67±47.92 | 19.12±33.11 |
| | *CD146 (MCAM)* | 4.13±6.66 | 0.96±1.67 |
| | *CD325* | 81.75±13.17 | 90.92±6.46 |

**Supplementary Table 1. Analysis of the surface antigen patterns of SC and DN preadipocytes.** Expression of 17 markers was determined in undifferentiated preadipocytes of 3 independent donors with T/C genotype for the rs1421085 locus in the *FTO* gene by flow cytometry. Four groups of markers were tested: hematopoietic/monocyte, endothelial, MSC/fibroblast markers, integrins and CAMs. The numbers represent the percentage of positive cells ±SD. SC: Subcutaneous, DN: Deep-neck, PA: preadipocyte.

**Supplementary Table 2.**

| GENE ID | ENCODED PROTEIN | ROLE IN ADIPOGENESIS |
|---|---|---|
| ***IGF1*** | Insulin-like growth factor I | Differentiation signal |
| ***SOX9*** | Transcription factor SOX-9 | Differentiation signal |
| ***PPARG*** | Peroxisome proliferator-activated receptor gamma | Adipogenic transcription factor |
| ***STAT3*** | Signal transducer and activator of transcription 3 | Differentiation signal |
| ***CEBPA*** | CCAAT/enhancer-binding protein alpha | Adipogenic transcription factor |
| ***SREBF1*** | Sterol regulatory element-binding protein 1 | Lipogenic gene regulators |
| ***SLC2A4*** | Glucose transporter member 4 | Adipocyte protein |
| ***FABP4*** | Fatty acid-binding protein. adipocyte | Adipocyte protein |
| ***LPL*** | Lipoprotein lipase | Adipocyte protein |
| ***AGPAT2*** | 1-acylglycerol-3-phosphate O-acyltransferase 2 | Adipocyte protein |
| ***PLIN1*** | Perilipin-1 | Adipocyte protein |
| ***ADIPOQ*** | Adiponectin | Adipocyte protein |
| ***LEP*** | Leptin | Adipocyte protein |
| ***STAT5A*** | Signal transducer and activator of transcription 5 | Adipogenesis regulator |
| ***EBF1*** | Transcription factor COE1/Early B-factor 1 | Adipogenesis regulator |
| ***CIDEC*** | Cell death inducing DFFA like effector C | Lipid droplet formation |
| ***PPARGC1A*** | Peroxisome proliferator-activated receptor gamma coactivator 1-alpha | Transcriptional co-factor |
| ***PPARGC1B*** | Peroxisome proliferator-activated receptor gamma coactivator 1-beta | Transcriptional co-factor |

**Supplementary Table 2. List of general adipocyte marker genes, the encoded proteins and their role in adipogenesis**

**Supplementary Table 3.**

| Genes | Encoded Protein | FC DNW_SCW/DNB_SCB | pValue DNW_SCW/DNB_SCB | FC DNB_DNW/SCB_SCW | pValue DNB_DNW/SCB_SCW |
|---|---|---|---|---|---|
| GROUP 1 | | | | | |
| *AC007938.2* | | -/2.6 | -/0.02 | 2.6/- | 0.03/- |
| *AC009126.1* | | -/1.8 | -/0.009 | 2.3/- | 0.0003/- |
| *AC034102.6* | | -/2.3 | -/0.01 | 2.3/- | 0.02/- |
| *AL606534.4* | | 3.6/3.2 | 0.003/0.005 | 2.7/2.9 | 0.04/0.03 |
| *ARNTL2* | Aryl hydrocarbon receptor nuclear translocator-like protein 2 | 2.1/2.3 | 0.001/4.2E-05 | 2.2/1.9 | 9.5E-04/1.5E-02 |
| *B3GNT5* | Lactosylceramide 1,3-N-acetyl-beta-D-glucosaminyltransferase | 2.2/1.9 | 0.009/0.04 | 2.2/2.6 | 0.009/0.002 |
| *CIDEA* | Cell death activator CIDE-A | 3.6/- | 0.03/- | 3.8/7.9 | 0.02/0.0001 |
| *CKMT1B* | Creatine kinase 1B | 3.1/- | | -/2.6 | -/0.04 |
| *CPA2* | Carboxypeptidase A2 | 2.9/- | 0.01/- | 3.5/4.9 | 0.0004/8.5E-05 |
| *CPT1B* | Carnitine O-palmitoyltransferase 1, muscle isoform | 1.8/2.1 | 0.015/0.0001 | 2.3/2.0 | 4.6E-05/8.2E-03 |
| *DAZL* | Deleted in azoospermia-like | -/3.9 | -/0.003 | 3.6/- | 0.01/- |
| *DRP2* | Dystrophin-related protein 2 | -/2.3 | -/0.007 | 2.4/- | 0.005/- |
| *FAM151A* | Protein FAM151A | 4.5/2.9 | 1.2E-05/0.002 | 2.4/3.7 | 0.03/0.0007 |
| *FAM189A2* | Protein FAM189A2 | -/2.6 | -/0.006 | 2.3/- | 0.03/- |
| *GPR160* | Probable G-protein coupled receptor 160 | -/2.5 | -/0.0003 | 2.4/- | 0.001/- |
| *KCNIP2* | Kv channel-interacting protein 2 | /2.4 | /0.02 | 2.7/2.3 | 0.008/0.03 |
| *KCNIP2-AS1* | | -/2.5 | -/0.01 | 2.5/- | 0.01/- |
| *LINC01347* | | 3.1/2.7 | 0.0007/0.01 | 2.6 | 0.05 |
| *LINC02458* | | -/2.6 | -/0.03 | 2.9/- | 0.01/- |
| *OASL* | 2'-5'-oligoadenylate synthase-like protein | 4.1/- | 0.006/- | 5.2/11.0 | 0.001/1.0E-06 |
| *PCDH19* | | 2.7/4.7 | 0.05/0.0001 | 4.4/- | 0.0005/- |
| *PM20D1* | N-fatty-acyl-amino acid synthase | 4.5/4.2 | 0.003/0.006 | 7.0/7.5 | 9.8E-05/5.8E-05 |
| *RASD1* | Dexamethasone-induced Ras-related protein 1 | 2.04/- | 0.02/- | 2.6/3.4 | 1.7E-03/2.6E-05 |
| *RASSF6* | Ras association domain-containing protein 6 | -/2.7 | -/0.03 | 2.9/- | 0.04/- |
| *RGL3* | Ral guanine nucleotide dissociation stimulator-like 3 | 2,4/- | 0.01/- | 2.3/3.1 | 2.8E-02/1.5E-03 |
| *RXRG* | Retinoic acid receptor RXR-gamma | 2.7/- | 0.02/- | -/2.8 | -/0.02 |
| *SCN4A* | Sodium channel protein type 4 subunit alpha | 3.7/- | 2.3E-05/- | 2.8/6.3 | 1.3E-03/5.7E-09 |
| *SLC19A3* | Thiamine transporter 2 | 1.9/- | 0.028/- | 2.0/2.1 | 4.6E-02/2.2E-02 |
| *SLC7A10* | Amino acids transporter | 5.3/4.3 | 2.6E-05/0.003 | 2.7/3.3 | 5.0E-02/1.6E-02 |

| | | | | | |
|---|---|---|---|---|---|
| GROUP 2 | | | | | |
| ***CPA4*** | Carboxypeptidase A4 | 2.4/2.1 | 0.03/0.04 | 4.8/5.3 | 1.4E-06/1.1E-06 |
| ***CX3CL1*** | Fractalkine | 4.8/- | 0.004/- | -/3.7 | -/4.0E-02 |
| ***CXCL16*** | C-X-C motif chemokine 16 | 1.9/- | 0.0008/- | -/1.9 | -/3.0E-03 |
| ***EGFLAM*** | Pikachurin | 2.1/- | 0.0004 | 1.9/2.4 | 0.009/0.0001 |
| ***EPB41L4B*** | Band 4.1-like protein 4B | 2.1/- | 0.02/- | 2.3/2.6 | 0.01/0.002 |
| ***LINC00623*** | | 2.4/- | 0.02/- | 2.6 | 2.0E-02 |
| ***MEST*** | Mesoderm-specific transcript homolog protein | 2.6/2.2 | 1.3E-05/0.0009 | 2.5/3.0 | 0.0001/2.6E-06 |
| ***MYOM1*** | Myomesin-1 | 2.0/- | 0.004/- | 1.9 | 1.0E-02 |
| ***SLC22A11*** | Solute carrier family 22 member 11 | -/3.7 | -/0.01 | 5.6/- | 0.002/- |
| ***SLCO2B1*** | organic anion transporter family member 2B1 | -/2.5 | -/0.03 | 3.1/- | 0.006/- |
| ***TSLP*** | Thymic stromal lymphopoietin | 2.5/2.3 | 0.002/0.002 | 2.5/2.7 | 0.0006/0.0007 |

**Supplementary Table 3. Common genes between anatomical location comparison and upregulated after brown differentiation protocol,** FC: fold change

**Supplementary Table 4.**

| Genes | Encoded Protein | FC DNW_SCW/DNB_SCB | pValue DNW_SCW/DNB_SCB | FC DNW_DNB/SCW_SCB | pValue DNW_DNB/SCW_SCB |
|---|---|---|---|---|---|
| GROUP 3 | | | | | |
| ***AL049825.1*** | | 3.6 | 0.04 | 4.6/- | 0.01/- |
| ***ANO4*** | Anoctamin-4 | 2.7/- | 0.006/- | 2.3 | 0.04 |
| ***CCDC146*** | Coiled-coil domain-containing protein 146 | 1.9/- | 2.6E-05/- | 1.8 | 0.0005 |
| ***CCNA1*** | Cyclin-A1 | 3.4 | 0.02 | 5.0/- | 0.002/- |
| ***DIO2*** | Type II iodothyronine deiodinase | 3.5/- | 0.004/- | 2.8 | 0.04 |
| ***FBN2*** | Fibrillin-2 | 5.9/6.7 | 3.9E-07/6.1E-08 | 2.6/- | 0.03/- |
| ***GPC3*** | Glypican-3 | 2.9/- | 0.0002/- | 2.9 | 0.001 |
| ***GPM6B*** | Neuronal membrane glycoprotein M6-b | 3.5/- | 5.3E-05/- | 3.2 | 0.0005 |
| ***IQCH-AS1*** | | 1.9/- | 0.002/- | 1.8 | 0.01 |
| ***KLF5*** | Krueppel-like factor 5 | 3.1/2.3 | 2.0E-05/0.005 | 1.9 | 0.04 |
| ***LINC00242*** | | 2.1/- | 0.04/- | 3.2 | 0.0009 |
| ***LINC00702*** | | 2.6/3.3 | 0.001/0.0002 | 3.9/2.9 | 0.0003 |
| ***LINC01238*** | | 5.6/- | 0.0007/- | 4.9 | 0.005 |
| ***LINC02268*** | | 5.8 | 0.009 | 2.6 | 0.05 |
| ***MARCO*** | Macrophage receptor MARCO | 3.6/- | 0.005/- | 3.1 | 0.03 |
| ***MOB3B*** | MOB kinase activator 3B | 2.6/2.1 | 0.001/0.03 | 2.2 | 0.02 |
| ***PPIAP39*** | | 2.7 | 0.005 | 3.4/2.4 | 0.0007/0.02 |
| ***ROBO2*** | Roundabout homolog 2 | 2.6/- | 0.02/- | 4 | 0.0005 |
| ***SLITRK1*** | SLIT and NTRK-like protein 1 | 6.5/6.3 | 2.7E-05/0.0002 | 3.4 | 0.02 |
| ***SSPO*** | SCO-spondin | 2.6/2.4 | 6E-05/0.001 | 1.8 | 0.04 |
| ***TRIM55*** | Tripartite motif-containing protein 55 | 2.9/- | 0.02/- | 3.3 | 0.02 |
| ***TRNP1*** | TMF-regulated nuclear protein 1 | 1.9/2.0 | 0.02/0.01 | 2.6/2.4 | 0.0003/0.0009 |
| ***TRPV6*** | Transient receptor potential cation channel subfamily V6 | 3.0/- | 0.03/- | 5 | 0.003 |
| ***ZNF385B*** | Zinc finger protein 385B | 3.2 | 0.04 | 6.7/3.3 | 0.0002/0.02 |
| GROUP 4 | | | | | |
| ***AL096865.1*** | | 4.5/9.2 | 0.0004/0.0002 | 4.9/- | 0.040 |
| ***COL8A1*** | Collagen alpha-1(VIII) chain | 3.4/2.8 | 0.0003/0.005 | /2.5 | 0.030 |
| ***CRISPLD2*** | Cysteine-rich secretory protein LCCL domain-containing 2 | 2.5 | 0.04 | /3.7 | 0.001 |
| ***DPT*** | Dermatopontin | 2.4/2.5 | 0.01/0.01 | 2.6/2.6 | 0.012/0.01 |
| ***GRIA1*** | Glutamate receptor 1 | 3.5 | 0.04 | /4.4 | 1.5E-02 |
| ***NEGR1*** | Neuronal growth regulator 1 | 1.8/- | 0.02/- | 2.9/2.7 | 5E-05/1.1E-05 |
| ***RSPO1*** | R-spondin-1 | 4.6/6.4 | 2.7E-05/0.2E-06 | 2.6/3.5 | 0.006/0.04 |
| ***TNC*** | Tenascin | 2.8/3.9 | 0.03/0.002 | /3.3 | 0.02/- |
| ***TRPC6*** | Short transient receptor potential channel 6 | 19.7/7.05 | 5.0E-15/1.5E-05 | 4.3/- | 7.0E-04 |

| | | | | | |
|---|---|---|---|---|---|
| ***ZNF467*** | Zinc finger protein 467 | 2.3/2.3 | 0.009/0.02 | 2.1 | 4.0E-02 |

**Supplementary Table 4. Common genes between anatomical location comparison and downregulated after brown differentiation protocol,** FC: fold change

#### Supplementary Table 5A.

| KEGG enriched pathways in FTO obese LOW expressed genes | KEGG enriched pathways in BATLAS | KEGG enriched pathways in ProFAT |
|---|---|---|
| ***Metabolic pathways*** | **Metabolic pathways** | *Metabolic pathways* |
| ***Fatty acid metabolism*** | **Thermogenesis** | *Parkinson's disease* |
| ***Thermogenesis*** | **Oxidative phosphorylation** | *Huntington's disease* |
| ***Oxidative phosphorylation*** | **Parkinson's disease** | *Thermogenesis* |
| ***Carbon metabolism*** | **Huntington's disease** | *Non-alcoholic fatty liver disease (NAFLD)* |
| ***Huntington's disease*** | **Non-alcoholic fatty liver disease (NAFLD)** | *Carbon metabolism* |
| ***Parkinson's disease*** | **Alzheimer's disease** | *Alzheimer's disease* |
| ***Fatty acid degradation*** | **Carbon metabolism** | *Oxidative phosphorylation* |
| ***Alzheimer's disease*** | **Citrate cycle (TCA cycle)** | *Citrate cycle (TCA cycle)* |
| ***Non-alcoholic fatty liver disease (NAFLD)*** | **Fatty acid degradation** | *2-Oxocarboxylic acid metabolism* |
| ***Valine. leucine and isoleucine degradation*** | **Fatty acid metabolism** | *Fatty acid metabolism* |
| ***Citrate cycle (TCA cycle)*** | **Valine. leucine and isoleucine degradation** | *Glyoxylate and dicarboxylate metabolism* |
| ***PPAR signaling pathway*** | **Cardiac muscle contraction** | *Fatty acid degradation* |
| ***Fatty acid elongation*** | **PPAR signaling pathway** | *Valine. leucine and isoleucine degradation* |
| ***Pyruvate metabolism*** | **2-Oxocarboxylic acid metabolism** | *Glycolysis / Gluconeogenesis* |
| ***Cardiac muscle contraction*** | **Pyruvate metabolism** | *Fatty acid elongation* |
| ***Glyoxylate and dicarboxylate metabolism*** | **Glyoxylate and dicarboxylate metabolism** | *Cardiac muscle contraction* |
| ***Glycolysis / Gluconeogenesis*** | **Glycolysis / Gluconeogenesis** | *Pyruvate metabolism* |
| ***2-Oxocarboxylic acid metabolism*** | **Fatty acid elongation** | *PPAR signaling pathway* |
| **Propanoate metabolism** | **Propanoate metabolism** | Biosynthesis of amino acids |
| **Peroxisome** | **Butanoate metabolism** | Retrograde endocannabinoid signaling |
| **Butanoate metabolism** | **Tryptophan metabolism** | Central carbon metabolism in cancer |
| **Adipocytokine signaling pathway** | **Peroxisome** | Glycine. serine and threonine metabolism |
| **Tryptophan metabolism** | **beta-Alanine metabolism** | |
| **beta-Alanine metabolism** | **Adipocytokine signaling pathway** | |
| Biosynthesis of unsaturated fatty acids | Retrograde endocannabinoid signaling | |
| Type I diabetes mellitus | Lysine degradation | |
| AMPK signaling pathway | Biosynthesis of amino acids | |
| Glucagon signaling pathway | Ribosome | |
| Insulin resistance | | |
| Allograft rejection | | |
| Steroid hormone biosynthesis | | |
| Graft-versus-host disease | | |

**Supplementary Table 5B.**

| REACTOM enriched pathways in FTO obese LOW expressed genes | REACTOM enriched pathways in BATLAS | REACTOM enriched pathways in ProFAT |
|---|---|---|
| ***Metabolism*** | **The citric acid (TCA) cycle and respiratory electron transport** | *The citric acid (TCA) cycle and respiratory electron transport* |
| ***Metabolism of lipids*** | **Metabolism** | *Metabolism* |
| ***The citric acid (TCA) cycle and respiratory electron transport*** | **Respiratory electron transport** | *Respiratory electron transport, ATP synthesis by chemiosmotic coupling, and heat production by uncoupling proteins.* |
| ***Fatty acid metabolism*** | **Respiratory electron transport, ATP synthesis by chemiosmotic coupling, and heat production by uncoupling proteins.** | *Respiratory electron transport* |
| ***Respiratory electron transport, ATP synthesis by chemiosmotic coupling, and heat production by uncoupling proteins.*** | **Pyruvate metabolism and Citric Acid (TCA) cycle** | *Pyruvate metabolism and Citric Acid (TCA) cycle* |
| ***Respiratory electron transport*** | **Citric acid cycle (TCA cycle)** | *Citric acid cycle (TCA cycle)* |
| ***Mitochondrial Fatty Acid Beta-Oxidation*** | **Fatty acid metabolism** | *Mitochondrial Fatty Acid Beta-Oxidation* |
| ***Pyruvate metabolism and Citric Acid (TCA) cycle*** | **Mitochondrial Fatty Acid Beta-Oxidation** | *Fatty acid metabolism* |
| ***Signaling by Retinoic Acid*** | **Glyoxylate metabolism and glycine degradation** | *Glyoxylate metabolism and glycine degradation* |
| ***mitochondrial fatty acid beta-oxidation of saturated fatty acids*** | **Metabolism of lipids** | *mitochondrial fatty acid beta-oxidation of saturated fatty acids* |
| ***Beta oxidation of hexanoyl-CoA to butanoyl-CoA*** | **mitochondrial fatty acid beta-oxidation of saturated fatty acids** | *Regulation of pyruvate dehydrogenase (PDH) complex* |
| ***Beta oxidation of decanoyl-CoA to octanoyl-CoA-CoA*** | **Beta oxidation of hexanoyl-CoA to butanoyl-CoA** | *Beta oxidation of hexanoyl-CoA to butanoyl-CoA* |
| ***Regulation of pyruvate dehydrogenase (PDH) complex*** | **Beta oxidation of decanoyl-CoA to octanoyl-CoA-CoA** | *Beta oxidation of decanoyl-CoA to octanoyl-CoA-CoA* |
| ***Glyoxylate metabolism and glycine degradation*** | **Signaling by Retinoic Acid** | *Signaling by Retinoic Acid* |
| ***Citric acid cycle (TCA cycle)*** | **Regulation of pyruvate dehydrogenase (PDH) complex** | *Metabolism of lipids* |
| *Import of palmitoyl-CoA into the mitochondrial matrix* | **Protein localization** | *Import of palmitoyl-CoA into the mitochondrial matrix* |
| **Pyruvate metabolism** | **Mitochondrial biogenesis** | Complex I biogenesis |
| **Beta oxidation of octanoyl-CoA to hexanoyl-CoA** | **Cristae formation** | Mitochondrial protein import |
| **Mitochondrial biogenesis** | **Beta oxidation of palmitoyl-CoA to myristoyl-CoA** | Transcriptional activation of mitochondrial biogenesis |
| **Beta oxidation of palmitoyl-CoA to myristoyl-CoA** | **Beta oxidation of lauroyl-CoA to decanoyl-CoA-CoA** | Lysine catabolism |
| **Beta oxidation of lauroyl-CoA to decanoyl-CoA-CoA** | **Beta oxidation of octanoyl-CoA to hexanoyl-CoA** | Branched-chain amino acid catabolism |
| **Cristae formation** | **Pyruvate metabolism** | Metabolism of amino acids and derivatives |
| **Peroxisomal protein import** | **Peroxisomal protein import** | |
| **Peroxisomal lipid metabolism** | **Peroxisomal lipid metabolism** | |
| **Protein localization** | Complex I biogenesis | |
| Endosomal/Vacuolar pathway | Mitochondrial translation initiation | |
| Signaling by Nuclear Receptors | Mitochondrial translation elongation | |
| Triglyceride metabolism | Mitochondrial translation termination | |
| mitochondrial fatty acid beta-oxidation of unsaturated fatty acids | Mitochondrial protein import | |

| | |
|---|---|
| Synthesis of very long-chain fatty acyl-CoAs | Lysine catabolism |
| Antigen Presentation: Folding, assembly and peptide loading of class I MHC | Branched-chain amino acid catabolism |
| Glycerophospholipid biosynthesis | Metabolism of amino acids and derivatives |
| Synthesis of bile acids and bile salts via 24-hydroxycholesterol | Beta oxidation of butanoyl-CoA to acetyl-CoA |
| Synthesis of bile acids and bile salts via 27-hydroxycholesterol | Gluconeogenesis |
| Interferon gamma signaling | Mitochondrial calcium ion transport |
| Acyl chain remodeling of CL | |
| Formation of ATP by chemiosmotic coupling | |
| Metabolism of steroids | |
| Linoleic acid (LA) metabolism | |
| RA biosynthesis pathway | |
| Synthesis of bile acids and bile salts via 7alpha-hydroxycholesterol | |
| TP53 Regulates Metabolic Genes | |
| Triglyceride catabolism | |
| PPARA activates gene expression | |
| Retinoid metabolism and transport | |
| alpha-linolenic acid (ALA) metabolism | |
| Antigen processing-Cross presentation | |
| Phospholipid metabolism | |
| Visual phototransduction | |
| Triglyceride biosynthesis | |
| Beta oxidation of myristoyl-CoA to lauroyl-CoA | |
| Immunoregulatory interactions between a Lymphoid and a non-Lymphoid cell | |

**Supplementary Table 5. Tables show the significantly enriched pathways identified by genes expressed lower in *FTO* C/C samples, BATLAS and ProFAT gene set. (A) KEGG pathways (B) REACTOM pathways.** Red letters: Pathways presented in all three samples; Bold: Common Pathways in *FTO* obesity-risk based DEGs and BATLAS; Italic font: Common Pathways in *FTO* obesity-risk based DEGs and ProFAT. Blue letters: Common Pathways only in BATLAS and ProFAT.

**Supplementary Table 6A**.

| Gene | BC score | Bridges | FC | Gene Description |
|---|---|---|---|---|
| ***ATP5B*** | 2015 | 18 | 2.2 | ATP synthase beta polypeptide |
| ***SDHB*** | 1028 | 18 | 2.0 | succinate dehydrogenase complex |
| ***SUCLG1*** | 1064 | 16 | 2.1 | succinate-CoA ligase |
| ***SOD2*** | 2418 | 14 | 2.4 | superoxide dismutase 2 |
| ***DECR1*** | 7717 | 13 | 1.8 | 2,4-dienoyl CoA reductase 1 |
| ***CD36*** | 1730 | 13 | 2.4 | Thrombospondin Receptor |
| ***ACADM*** | 1531 | 13 | 2.2 | acyl-CoA dehydrogenase |
| ***NDUFS1*** | 200 | 13 | 1.8 | NADH dehydrogenase Fe-S protein 1 |
| ***MDH1*** | 968 | 12 | 2.7 | malate dehydrogenase 1 |
| ***PPARGC1A*** | 854 | 12 | 2.6 | peroxisome proliferator-activated receptor gamma |
| ***UQCRC2*** | 139 | 12 | 2.0 | ubiquinol-cytochrome c reductase core protein II |
| ***ACSL1*** | 525 | 11 | 1.8 | acyl-CoA synthetase long-chain family member 1 |
| ***CPT1B*** | 294 | 11 | 2.0 | carnitine palmitoyltransferase 1B |
| ***CCL2*** | 4845 | 10 | 2.5 | chemokine (C-C motif) ligand 2 |
| ***LIPE*** | 3792 | 10 | 2.6 | lipase hormon sensitive |

**Supplementary Table 6B.**

| Gene | BC score | Bridges | FC | Gene Description |
|---|---|---|---|---|
| ***FN1*** | 17619 | 37 | 2.1 | fibronectin 1 |
| ***CTGF*** | 5508 | 16 | 2.6 | connective tissue growth factor |
| ***VCAN*** | 1117 | 16 | 2.4 | versican |
| ***TGFB1*** | 5194 | 14 | 1.9 | transforming growth factor, beta 1 |
| ***COL1A1*** | 4343 | 14 | 2.7 | collagen, type I, alpha 1 |
| ***CDH2*** | 3971 | 14 | 2.6 | cadherin 2, type 1, N-cadherin (neuronal) |
| ***BDNF*** | 5515 | 13 | 2.0 | brain-derived neurotrophic factor |
| ***VCAM1*** | 3666 | 12 | 3.1 | vascular cell adhesion molecule 1 |
| ***TIMP1*** | 2592 | 12 | 2.5 | TIMP metallopeptidase inhibitor 1 |
| ***LOX*** | 1377 | 12 | 2.0 | lysyl oxidase |
| ***VEGFC*** | 3627 | 11 | 1.9 | vascular endothelial growth factor C |
| ***FOS*** | 2986 | 9 | 2.0 | FBJ murine osteosarcoma viral oncogene homolog |
| ***SDC1*** | 2729 | 9 | 2.4 | syndecan 1 |
| ***BGN*** | 1644 | 9 | 3.9 | biglycan |
| ***GLI1*** | 5192 | 8 | 2.7 | GLI family zinc finger 1 |

**SupplementaryTable 6. Betweenness centrality (BC) score and Bridges number identified based on STRING Interactome network and Fold Changes (FC) identified by DESeq R of the differentially expressed genes based on the presence of the FTO obesity risk allele.** Analyses of the lower expressed genes **(A)** and the higher expressed genes **(B)** in FTO obesity-risk samples, n=6 donors

**Supplementary Table 7.**

| Genes less expressed in FTO C/C and high in DN samples | FC | TRANSCRIPT |
|---|---|---|
| ***CKMT1A*** | 4.9 | creatine kinase, mitochondrial 1A |
| ***LINC01348*** | 4.6 | Long Intergenic Non-Protein Coding RNA, nearest gene TOMM20 |
| ***PRSS35*** | 4.3 | Protease, serine35 |
| ***SPTA1*** | 3.6 | Spectrin, alpha, erythrocytic 1 (elliptocytosis 2) |
| ***EPB42*** | 3.5 | erythrocyte membrane protein band 4.2 |
| ***ZDHHC19*** | 3.2 | zinc finger, DHHC-type containing 19 |
| ***SNCG*** | 3.1 | Synuclein, gamma (breast cancer-specific protein 1) |
| ***HYAL1*** | 3.0 | hyaluronoglucosaminidase 1 |
| ***EMB*** | 3.0 | embigin |
| ***LINC02458*** | 2.8 | Long Intergenic Non-Protein Coding RNA, nearest gene AC006199.1 pseudogene |
| ***CKMT1B*** | 2.8 | creatine kinase, mitochondrial 1B |
| ***GBP1P1*** | 2.8 | guanylate binding protein 1, interferon-inducible pseudogene 1 |
| ***LINC01914*** | 2.8 | Long Intergenic Non-Protein Coding RNA, nearest gene c2orf91 |
| ***RHOXF1-AS1*** | 2.7 | Rhox Homebox family Member 1 antisense RNA |
| ***SGK2*** | 2.5 | serum/glucocorticoid regulated kinase 2 |
| ***FABP3*** | 2.5 | fatty acid binding protein 3, muscle and heart (mammary-derived growth inhibitor) |
| ***C1QTNF1*** | 2.5 | C1q and tumor necrosis factor related protein 1 |
| ***RPS6KL1*** | 2.5 | ribosomal protein S6 kinase-like 1 |
| ***AKR1C8P*** | 2.5 | Aldo-Keto Reductase Family 1 Member C8, Pseudogene |
| ***SOD2*** | 2.4 | superoxide dismutase 2, mitochondrial |
| ***PLA2G4A*** | 2.4 | phospholipase A2, group IVA (cytosolic. calcium-dependent) |
| ***FAM189A2*** | 2.3 | family with sequence similarity 189, member A2 |
| ***GPLD1*** | 2.2 | glycosylphosphatidylinositol specific phospholipase D1 |
| ***SLC19A3*** | 2.1 | solute carrier family 19 (thiamine transporter), member 3 |
| ***CPT1B*** | 2.0 | carnitine palmitoyltransferase 1B (muscle) |
| ***TPRG1*** | 2.0 | tumor protein p63 regulated 1 |
| ***LINC01140*** | 2.0 | LINC- RNA, complement to AC093155.3 protein coding transcript, Long Intergenic Non-Protein Coding RNA |
| ***RNF207*** | 1.9 | ring finger protein 207 |
| ***AC010319.5*** | 1.9 | LINC-RNA, complement to AC010319.2 protein coding transcript, nearest genes TMEM221. NXNL1 |
| ***GCKR*** | 1.9 | glucokinase (hexokinase 4) regulator |
| ***EHHADH*** | 1.9 | enoyl-CoA, hydratase/3-hydroxyacyl CoA dehydrogenase |
| ***AKAP1*** | 1.9 | A kinase (PRKA) anchor protein 1 |
| ***CRYAB*** | 1.8 | crystallin. alpha B |

**Supplementary Table 7. Genes that are significantly less expressed in the FTO obesity-risk genotypes (C/C) as compared to normal (T/T) and higher in DN as compared to SC samples.** FC: fold change after white and brown differentiation, average fold change (FC) when two data considered.

**Supplementary Table 8.**

| **Genes highly expressed in FTO C/C and low in DN samples** | **FC** | **TRANSCRIPT** |
|---|---|---|
| ***HAPLN1*** | 8.9 | hyaluronan and proteoglycan link protein 1 |
| ***CMKLR1*** | 7.8 | chemokine-like receptor 1 |
| ***CADPS*** | 6.0 | Ca++-dependent secretion activator |
| ***PTPRD-AS1*** | 5.4 | Protein Tyrosine Phosphatase Receptor Type D antisense 1 |
| ***IL11*** | 5.0 | interleukin 11 |
| ***COL13A1*** | 4.7 | Collagen, type XIII. alpha 1 |
| ***ACAN*** | 4.7 | aggrecan |
| ***AC244153.1*** | 4.5 | novel transcript, lncRNA antisense to ARHGAP23 (Rho GTPase Activating Protein 23) |
| ***COL8A2*** | 4.4 | Collagen, type VIII. alpha 2 |
| ***OPCML*** | 4.2 | opioid binding protein/cell adhesion molecule-like |
| ***NRG1*** | 4.1 | neuregulin 1 |
| ***LINC00900*** | 4.1 | long intergenic non-protein coding RNA 900 |
| ***EGR2*** | 3.9 | early growth response 2 |
| ***F2RL2*** | 3.9 | coagulation factor II (thrombin) receptor-like 2 |
| ***TENM2*** | 3.9 | teneurin transmembrane protein 2 |
| ***PCDH10*** | 3.9 | protocadherin 10 |
| ***CLEC2L*** | 3.8 | C-type lectin domain family 2, member L |
| ***IL27RA*** | 3.8 | interleukin 27 receptor, alpha |
| ***COL8A1*** | 3.7 | Collagen, type VIII. alpha 1 |
| ***MYH1*** | 3.6 | Myosin, heavy chain 1, skeletal muscle, adult |
| ***ULBP1*** | 3.5 | UL16 binding protein 1 |
| ***AC114284.1*** | 3.4 | novel transcript, lncRNA |
| ***ITGA11*** | 3.3 | Integrin, alpha 11 |
| ***AC079336.5*** | 3.3 | Novel Transcript, Antisense MYO1D |
| ***GDNF*** | 3.1 | glial cell derived neurotrophic factor |
| ***CDH6*** | 3.0 | cadherin 6, type 2, K-cadherin (fetal kidney) |
| ***HS3ST3A1*** | 2.9 | heparan sulfate (glucosamine) 3-O-sulfotransferase 3A1 |
| ***CASS4*** | 2.9 | Cas scaffolding protein family member 4 |
| ***F2R*** | 2.9 | coagulation factor II (thrombin) receptor |
| ***GDF6*** | 2.9 | growth differentiation factor 6 |
| ***PODNL1*** | 2.9 | podocan-like 1 |
| ***HCN1*** | 2.8 | hyperpolarization activated cyclic nucleotide-gated potassium channel 1 |
| ***COL1A1*** | 2.7 | Collagen, type I. alpha 1 |
| ***EVA1A*** | 2.7 | eva-1 homolog A (C. elegans) |

| | | |
|---|---|---|
| ***SHANK1*** | 2.7 | SH3 and multiple ankyrin repeat domains 1 |
| ***EPHB3*** | 2.6 | EPH receptor B3 |
| ***PRR16*** | 2.6 | proline rich 16 |
| ***PRSS12*** | 2.6 | Protease, serine 12 (neurotrypsin, motopsin) |
| ***SEPT4*** | 2.5 | Septin 4 |
| ***HACD4*** | 2.4 | 3-Hydroxyacyl-CoA Dehydratase 4 |
| ***RASGRP2*** | 2.4 | RAS guanyl releasing protein 2 (calcium and DAG-regulated) |
| ***NCKAP5*** | 2.4 | NCK-associated protein 5 |
| ***STAC*** | 2.4 | SH3 and cysteine rich domain |
| ***SDC1*** | 2.4 | syndecan 1 |
| ***KCNG1*** | 2.3 | potassium voltage-gated channel, subfamily G. member 1 |
| ***GDNF-AS1*** | 2.3 | GDNF antisense RNA 1 (head to head) |
| ***CDH11*** | 2.3 | cadherin 11, type 2, OB-cadherin (osteoblast) |
| ***ALDH3B1*** | 2.3 | aldehyde dehydrogenase 3 family, member B1 |
| ***FOXL1*** | 2.3 | forkhead box L1 |
| ***HOXC8*** | 2.3 | homeobox C8 |
| ***FAM86GP*** | 2.2 | family with sequence similarity 86, member G. pseudogene |
| ***TBX5*** | 2.2 | T-box 5 |
| ***MYO1D*** | 2.2 | myosin ID |
| ***FZD7*** | 2.1 | frizzled family receptor 7 |
| ***TGFB1I1*** | 2.1 | transforming growth factor beta 1 induced transcript 1 |
| ***IL20RA*** | 2.1 | interleukin 20 receptor, alpha |
| ***MRC2*** | 2.1 | mannose receptor, C type 2 |
| ***FSCN1*** | 2.1 | fascin homolog 1, actin-bundling protein (Strongylocentrotus purpuratus) |
| ***ACVR2A*** | 2.1 | activin A receptor, type IIA |
| ***BAIAP2L1*** | 2.1 | BAI1-associated protein 2-like 1 |
| ***LMO7*** | 2.1 | LIM domain 7 |
| ***ASNS*** | 2.1 | asparagine synthetase (glutamine-hydrolyzing) |
| ***GALNT5*** | 2.1 | UDP-N-acetyl-alpha-D-galactosamine:polypeptide N-acetylgalactosaminyltransferase 5 (GalNAc-T5) |
| ***SLIT2*** | 2.1 | slit homolog 2 (Drosophila) |
| ***STARD4-AS1*** | 2.0 | StAR Related Lipid Transfer Domain Containing 4 antisense RNA 1 |
| ***IRX3*** | 1.9 | iroquois homeobox 3 |
| ***RUBCNL*** | 1.9 | Rubicon Like Autophagy Enhancer |
| ***TBX5-AS1*** | 1.9 | TBX5 antisense RNA 1 |
| ***SPON1*** | 1.8 | spondin 1, extracellular matrix protein |
| ***WNT5B*** | 1.8 | wingless-type MMTV integration site family, member 5B |

**Supplementary Table 8. Genes that have significantly higher expression in the FTO obesity-risk genotypes (C/C) as compared to normal (T/T) and low in DN as compared to SC samples.** FC: fold change after white and brown differentiation, average fold change (FC) when two data considered
